# Reductive Analytics on Big MS Data leads to tremendous reduction in time for peptide deduction

**DOI:** 10.1101/073064

**Authors:** Muaaz Gul Awan, Fahad Saeed

## Abstract

In this paper we present a feasibility of using a data-reductive strategy for analyzing big MS data. The proposed method utilizes our reduction algorithm MS-REDUCE and peptide deduction is accomplished using Tide with hiXcorr. Using this approach we were able to process 1 million spectra in under 3 hours. Our results showed that running peptide deduction with smaller amount of selected peaks made the computations much faster and scalable with increasing resolution of MS data. Quality assessment experiments performed on experimentally generated datasets showed good quality peptide matches can be made using the reduced datasets. We anticipate that the proteomics and systems biology community will widely adopt our reductive strategy due to its efficacy and reduced time for analysis.

## I Technical Brief

Modern day proteomics involves the use of Mass Spectrometry for correctly sequencing the proteins [1]. This method involves breaking down of proteins into smaller peptides which are then fragmented inside a Mass Spectrometer [2]. The resultant spectra are then sequenced using two popular approaches i.e. Denovo Sequencing [3] and Database search [4]. Over the years the Database search method has gained much importance and many approaches have emerged for efficiently deducing peptides from Mass Spectra. Some of the popular database search approaches are Sequest [4], Mascot [5] and X!tandem [6]. Recently with the advent of more powerful Mass Spectrometry instruments huge number of high resolution spectra can be generated per second [7] [8]. Processing such huge number of spectra is a time taking process especially when combined with increasing resolution of the data. As a possible solution, numerous techniques have been introduced for reducing the amount of this data. Some algorithms achieve this by discarding the prospective noisy or low confidence spectra [9] [10]. While others take the more popular approach of discarding the unwanted peaks [11] [12]. True advantage of these algorithms had not been utilized till now because of two possible reasons; i) the data/noise reducing algorithms were very slow and provided too much overhead and ii) the existing peptide deduction algorithms did not exploit the reduced number of peaks in each spectrum.

A reimplementation of Sequest known as Tide [13] performs the database search manifolds faster than the original Sequest. However when dealing with high resolution instruments Tide does not provide a scalable performance [8]. Tide distributes the peaks of a spectrum across several bins. The size of these bins is defined by user in accordance with the resolution of the Mass Spectrometry instrument used. A smaller bin size is used to accommodate a higher resolution instrument. Smaller bin size corresponds to larger number of bins. Number of bins increases exponentially with decreasing bin size.

Like Sequest, Tide makes use of cross correlation or *X_corr_* score for determining the similarity between an experimentally obtained spectrum and a theoretically generated one (from the database). For Tide time taken for calculating *X_corr_* is proportional to the number of bins [8] also most of time is spent on calculating *X_corr_*. This an exponential increase in number of bins with increasing resolution results in an equivalent increase in time, this is has been presented in [8]. Number of bins created are independent of number of peaks in a given spectra. As a consequence a spectrum from a higher resolution instrument with handful of peaks will result in creation of large number of bins hence a larger processing time. To solve this problem a solution was presented in the form of an algorithm called *hiXcorr* which is able to calculate the *X_corr_* score in a time proportional to the number of peaks in spectrum. It has been shown that the time taken for calculating *X_corr_* using hiXcorr does not increase significantly when run for higher resolutions[8]. Thus hiXcorr provides an opportunity to exploit the abilities of data reduction algorithms.

From the above discussion it becomes apparent that processing time of hiXcorr is bound to increase with increasing number of peaks. But as discussed in [11] about 90% of the peaks in a spectrum do not play any part in correct deduction of peptide. Removal of these unnecessary peaks will not only make the data more manageable but will also provide the much needed speed up while using hiXcorr algorithm for calculating *X_corr_* score. We have recently shown that MS-REDUCE [11] is able to reduce peaks in each spectrum by 70% and still give high quality results. Also the data reduction process by MS-REDUCE is performed in just a fraction of time of peptide deduction process while performing the task 100x faster than other similar tools. Thus it makes sense to use MS-REDUCE on the datasets before running them through peptide deduction process.

On the basis of above discussion we present a data reductive strategy for protein deduction, we suggest using MS-REDUCE in the protein deduction pipeline along with hiXcorr integrated Tide. As showing in Fig. 1, MS-REDUCE can be used as an integral part of the process. This leads to much faster and more scalable solution for peptide deduction. The biggest advantage of this approach is the scalability it provides for increasing resolution of modern devices. We performed the quailty assessment analysis on thirteen experimentally generated datasets and for scalablility analysis we ran our experiments for over 1 million spectra.

**Figure 1:**
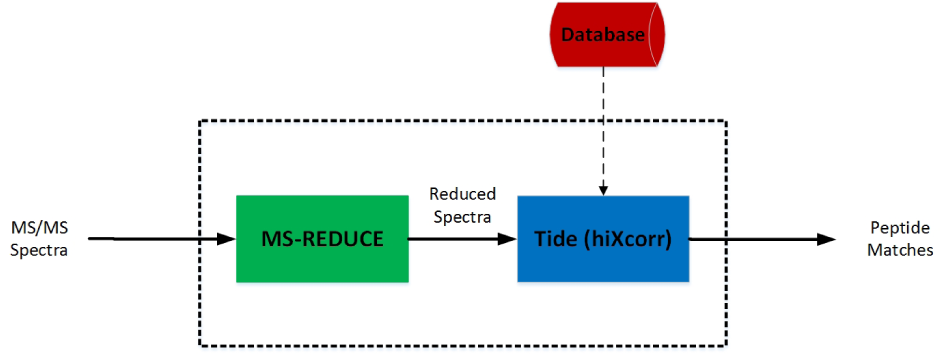
Figure showing the proposed data reductive strategy for peptide deduction.

We performed quality assessment analysis on thirteen experimentally genreated datasets generated using different fragmentation strategies i.e. HCD and CID. We used the same datasets in [11].

A piece of rat liver, which was freshly isolated was minced and then sonicated in guanidine-HCL(6M, 3ml). A peptide standard corresponding to the C-terminal sequence of the water channel Aquaporin-2 (AQP2) from rat, (BiotinLCCEPDTDWEEREVRRRQS*VELHS*PQSLPRGSKA) phosphorylated at both S256 and S261 were added to 500 g aliquots of liver sample (prior to trypsinization) and distinct amounts of 0.2 nmol, 20 pmol and 2 pmol were added. We will further reference these data sets as DS1, DS2 and DS3 respectively. We repeated the above procedure for AQP2 peptide standard (BiotinLCCEPDTDWEEREVRRRQSVELHSPQS*LPRGSKA) phosphorylated at S264, with amounts of 0.2 nmol, 20 pmol and 2 pmol.We will further reference these data sets as DS4, DS5 and DS6 respectively. These samples were then desalted, and were suspended in 0.1perc formic acid before being analyzed in Mass Spectrometer using HCD and CID fragmentation. Name of each dataset will be preceded by the type of fragmentation for example HCD-DS5 means DS5 with HCD fragmentation. Spectra obtained were then searched using Tide. A rat proteome database was obtained from (http://uniprot.org). Following static modifications were made while generating the idnexed database: carbamidomethyl (C: +57.021 Da) and dynamic modifications: Phospho (S,T,Y: +79.966), Deamidation (N,Q: +0.984), Oxidation (M: +15.995).

We also made use of Proteomics Dynamic Range Standard Set (UPS2) from Sigma-Aldrich. UPS2 data set contains a mixture of 48 individual human sequence recombinant proteins, each of which was selected to limit heterogeneous posttranslational modifications. It is formulated from 6 mixtures of 8 proteins to present a dynamic range of 5 orders of magnitude, ranging from 50pmol to 500amol. Briefly, 10.6 ug total protein (one vial) was resuspended in 50 ul of denaturation solution (8M urea, 50 mM TrisHCl, 75 mM NaCl) followed by reduction and alkylation. Samples were then treated with enzyme Trypsin at a 1:20 (w/w) ratio for 16 hours at 37C. Peptides obtained were then desalted (PepClean C-18 Spin Columns, Thermo Scientific) and eluted in 0.1perc formic acid. Sample amounts of 10, 50, and 200 ng of digested peptides were analyzed on an LTQ Orbitrap Velos (Thermo Scientific). Refseq Database (National Center for Biotechnology Information, March 3, 2010, 30,734 entries) was used for peptide searching. This database also contains the sequences for all human proteins included in the standard set UPS2, along with a list of common contaminating proteins. The indexed database was generated with following modifications: Carbamidomethylation of cysteine (+57.021 Da) as static modification and a variable modifications included oxidation of methionine (+15.995 Da) and deamidation of asparagine and glutamine (+0.984 Da). The UPS2 dataset was then appended multiple times to obtain a list of over 1 million spectra, this list of million spectra was then used for scalability study of this strategy.

Plots in Fig. 2 show a considerable speed up when using MS-REDUCE in conjunction with hiXcorr version of Tide. We performed experiments by varying the reduction ratio of MS-REDUCE and the bin-width used for Tide. It can be observed from the figure that for higher reduction factor i.e. when larger number of peaks are retained the processing time rises with increasing resolution. While for lower reduction factors it shows a very scalable performance. For 10% reduction factor the processing time almost becomes constant for larger resolution.

**Figure 2:**
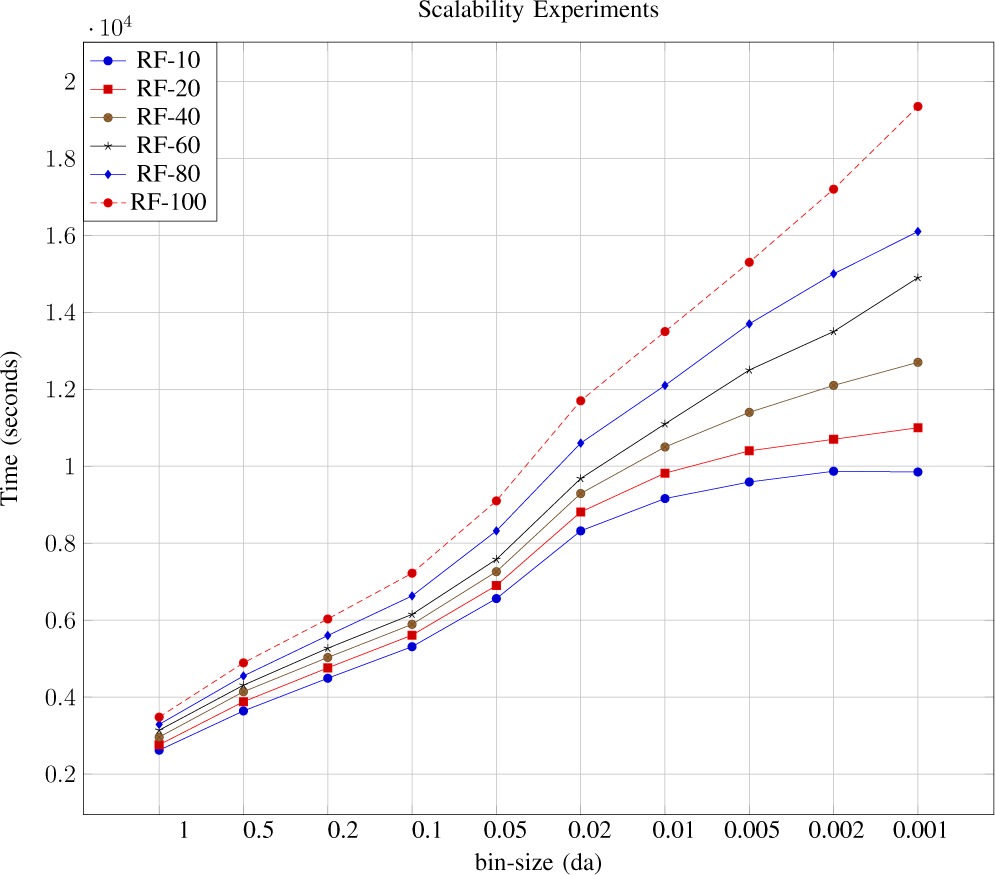
Figure showing timing plots of peptide deduction process using Tide with hiXcorr algorithm. Here RF represents Reduction Factor at which MS-REDUCE was run while treating the spectra. A decreasing Reduction Factor makes the process more scalable.

When the above experiments were repeated by using the original Tide software instead of Tide with hiXcorr, we observed near exponential rise in processing time with increasing resolution. It can be observed in Fig. 3, varying Reduction factor does not have any significant effect on the processing time. This is because original Tide calculates the *X_corr_* score in a time proportional to the number of bins and smaller bin size results in a very large number of bins for increasing resolution.

**Figure 3:**
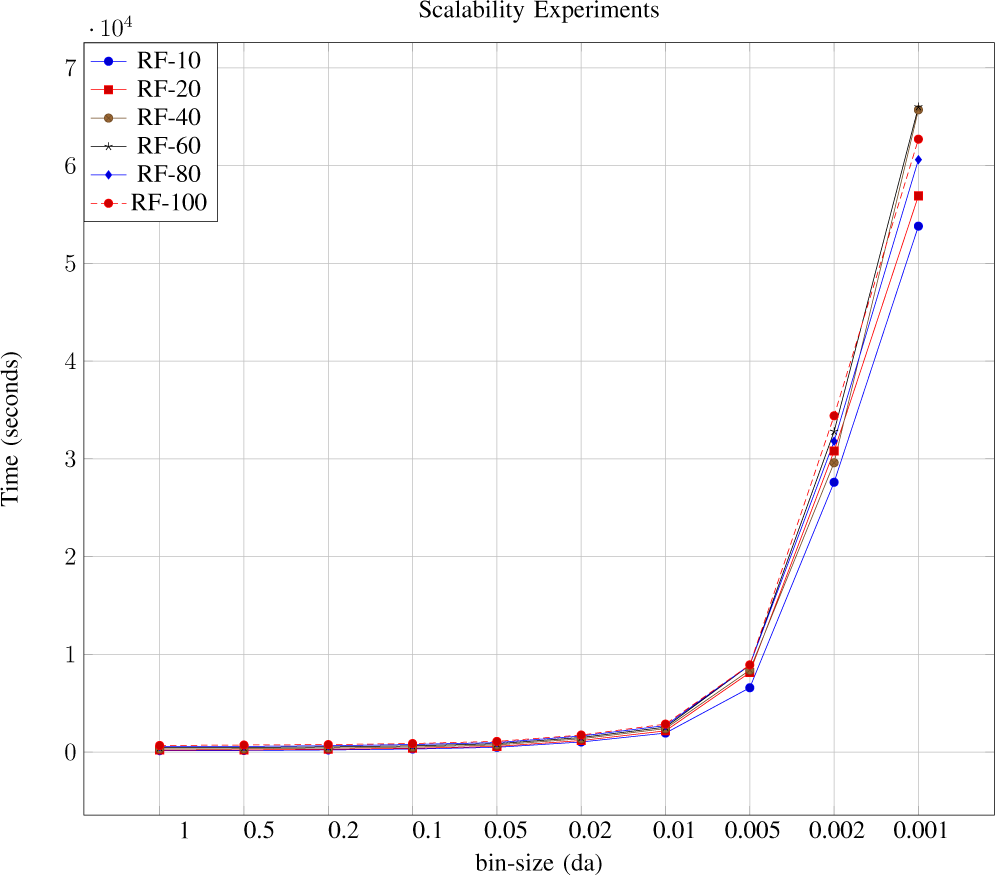
Figure showing timing plots of peptide deduction process using Tide with original *X_corr_* algorithm. Here RF represents Reduction Factor at which MS-REDUCE was run while treating the spectra. A decreasing Reduction Factor does not significantly affect the process.

For evaluating the effect of data reduction on the quality of peptide matches we performed quality assessment experiments using percolator [14] to post process the peptide spectral matches. We used the same method of quality assessment as used in [11]. The 4 shows even with reduced amount of data a large percentage of high quality matches was obtained. Thus a data reductive strategy can give sufficient high quality peptide matches while making the process much more scalable especially for modern high resolution instruments. We anticipate that reductive strategies such as MS-REDUCE [11] will gain popularity among proteomics tools developers due to its time-advantages. This will in turn lead to more scalable solutions even with high resolution MS data.

**Figure 4:**
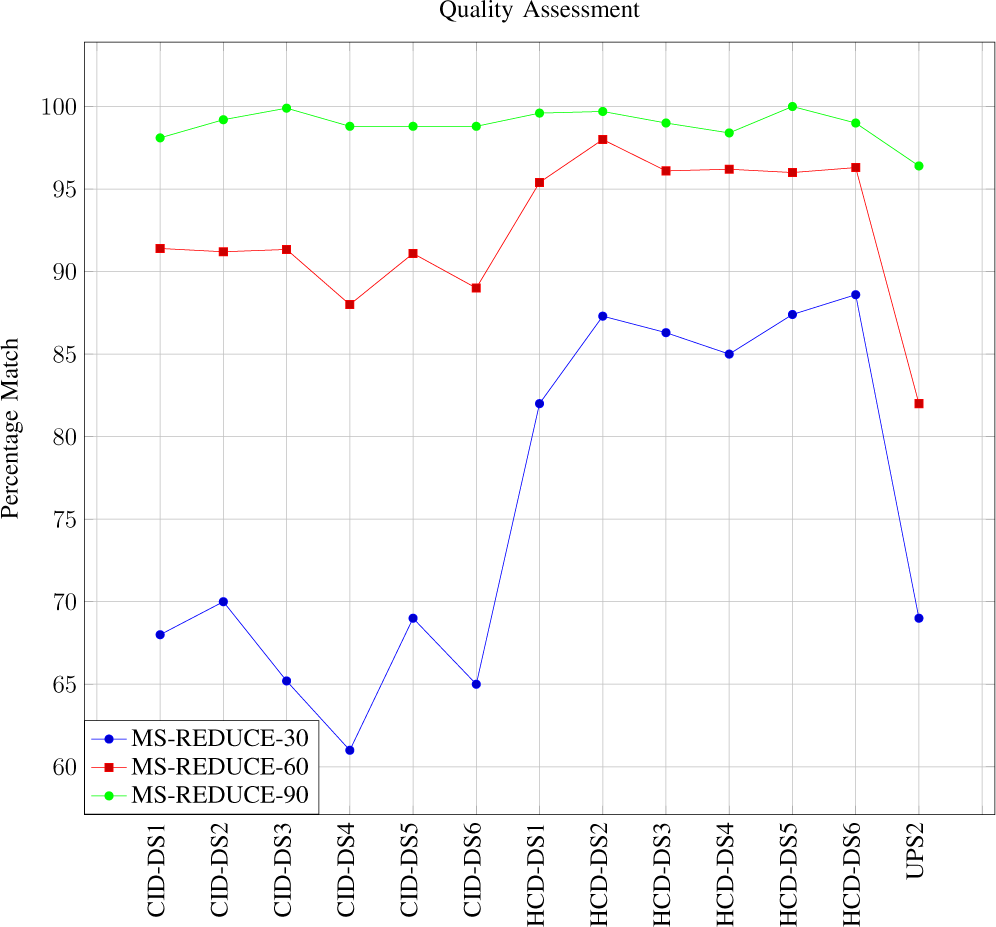
Figure showing quality assessment plots for MS-REDUCE. Quality Assessment plots for different Reduction Factor of MS-REDUCE can be observed. In the legend a numerical value with MS-REDUCE represents its reduction factor. X-axis contain the labels for the experimental datasets while Y-axis represents the percentage of peptide matches obtained from each dataset after being processed by each algorithm.

## II Acknowledgement

This work was supported in part by grants NSF CRII CCF-1464268.

